# Voxelotor (Oxbryta) Binds Multiple Hemoglobin Sites and Influences Protein Structure

**DOI:** 10.1101/2025.05.12.653546

**Authors:** Michael de la Cueva Tamanaha, Esmeralda Flores Cabrera, Joshua Sargeant, Paul D. Gershon, Premila P. Samuel Russell, Melanie J. Cocco

**Affiliations:** Department of Molecular Biology and Biochemistry, UC Irvine; Department of Chemistry, St. Louis University; Department of Pharmaceutical Sciences, UC Irvine

## Abstract

Voxelotor (Oxbryta, GBT440) is a first-in-class drug, FDA-approved to treat sickle cell disease in 2019 but withdrawn from market in 2024. This drug acts as an allosteric modulator, designed to shift the equilibrium to the oxygenated R conformation. The drug was shown to both limit the accumulation of deoxygenated T-conformation sickle cell Hb fibers and increase Hb oxygen affinity. X-ray crystallography previously showed one-to-one Voxelotor binding stoichiometry for Hb, with the drug molecule bound to N-terminus of an alpha subunit. Here we use NMR spectroscopy to assess the structure of Voxelotor-bound hemoglobin in solution and mass spectrometry (MS) to determine stoichiometry and sites of binding. We find that the structure and stoichiometry of binding are far more heterogeneous than previously described. The addition of Voxelotor to R-conformation Hb induces an NMR signal found in the T-conformation of Hb. In addition, MS shows that the drug binds Hb at multiple sites, including the N-terminus of the beta subunit. The properties of Hb with Voxelotor bound at secondary sites have not been explored but should be considered at high doses. Heterogeneous binding should be assessed in other drugs of this class including GBT(021)601 currently in clinical trial.

## Introduction

The approval of Voxelotor to treat sickle cell disease (SCD) was welcomed in 2019 as a needed oral therapy for a condition with severe and painful symptoms. This was the first drug to treat SCD by binding hemoglobin (Hb) directly; assessments of efficacy were very positive^1^. Voxelotor was shown to delay fiber formation *in vitro* and reduce sickling^2^. The drug is readily absorbed and partitions very efficiently into erythrocytes^3^. Voxelotor contains a reactive aldehyde that can form a Schiff base, a covalent, reversible adduct with primary amines. Complementary binding to the cleft between Hb α subunits enables the drug to target the N-terminus of one α subunit. The crystal structure of Voxelotor-bound Hb revealed a single drug molecule bound to the R-state. Voxelotor inhibits sickling by shifting the equilibrium away from the fiber-forming T-state in sickle cell Hb (HbSS)^1^. Unfortunately, use in humans showed significant adverse effects and the drug was withdrawn in 2024^4^.

We now know that Hb can exist within a continuum of conformations between static R and T states seen by crystallography^5^. High-resolution NMR measurements in solution have highlighted structural features not observed when Hb is constrained in solid-state crystal structures^6,7^. In addition, NMR has detected hybrid conformers of Hb with ligand bound (R-state) and T-state features^8,9^. Since Voxelotor is an allosteric regulator, we set out to understand its mechanism of action by studying the structure of Voxelotor-bound Hb in solution using NMR spectroscopy. Although the Voxelotor-bound Hb crystal structure was in the R-state conformation^1^, our NMR studies reveal that a spectral feature of the deoxy T-conformation appears upon drug binding in the presence of ligands that normally drive formation of the R-state (oxygen or carbon monoxide). Here we describe new structural information provided by solution-state NMR studies. Additionally, we find that the stoichiometry of binding is considerably more complex than the simple 1:1 interaction between drug and Hb tetramer previously assumed.

## Methods

### Detailed procedures are provided in Supplemental Information

Voxelotor was purchased from MedKoo or MedChem Express. Hemoglobin was purified from blood freshly donated for research purposes to the UC Irvine Experimental Tissue Repository. Blood was deidentified except for sickle cell status. HbSS lysates were prepared as previously described^10^. Normal HbA purification was based on the protocol of Sun and Palmer^11^. Carbonmonoxy HbA (HbACO) NMR samples were prepared in 5 mM sodium phosphate, then diluted to 100 mM sodium phosphate, pH 7.2 or pH 6.9, 10% D_2_O. Oxygenated HbA (HbAO_2_) samples for NMR experiments prepared in 5 mM sodium phosphate, pH 7.6, 10% D_2_O. NMR 1D and 2D NOESY (τ_mix_ = 100 ms) experiments were performed using a Bruker Neo 800 MHz spectrometer at 25 °C. Voxelotor has limited solubility in aqueous solution. Since we observed drug aggregation and turbidity, NMR samples were centrifuged prior to study and the amount of remaining drug bound to Hb was quantitated by LC/MS. 100 mM sodium borohydride was used to trap the Schiff base; Voxelotor was then quantitated using a Xevo G2-XS QTOF (Waters) LC/MS. Tryptic digests were analyzed using nanoLC MS/MS to determine sites of binding. Computational modeling of Voxelotor-bound to the Hb β subunit was accomplished using Schrödinger Release 2025-2^12^.

## Results and Discussion

### LC/MS Reveals Multiple Drug Binding Sites

Our biophysical studies were initiated by determining the stoichiometry of Voxelotor binding to Hb. The Schiff base formed between drug and protein is labile, especially under the acidic conditions typically used in LC/MS (or HPLC). Quantitation of bound drug requires reducing agents to trap the drug in the bound state. We replicated a published study of Voxelotor added to HbSS lysates^10^ with one notable difference (**Figure 1A**). We used a strong reducing agent, sodium borohydride, whereas the previous study used a weak reductant, sodium cyanoborohydride, known to be up to five times less effective in Schiff base trapping^13^. We confirmed the inefficiency of sodium cyanoborohydride in quantitating the amount of Hb-bound Voxelotor (**Figure S1**). Voxelotor added to HbSS lysates showed strong binding to the N-terminus of the α subunit as expected ^10^. Binding to the β N-terminus was previously reported but dismissed as unlikely since it was believed to occur only at very high concentrations of drug (10-fold excess)^10^. Here, we find new evidence of binding to the β subunit at concentrations achievable *in vivo*^3^.

**Figure 1.**
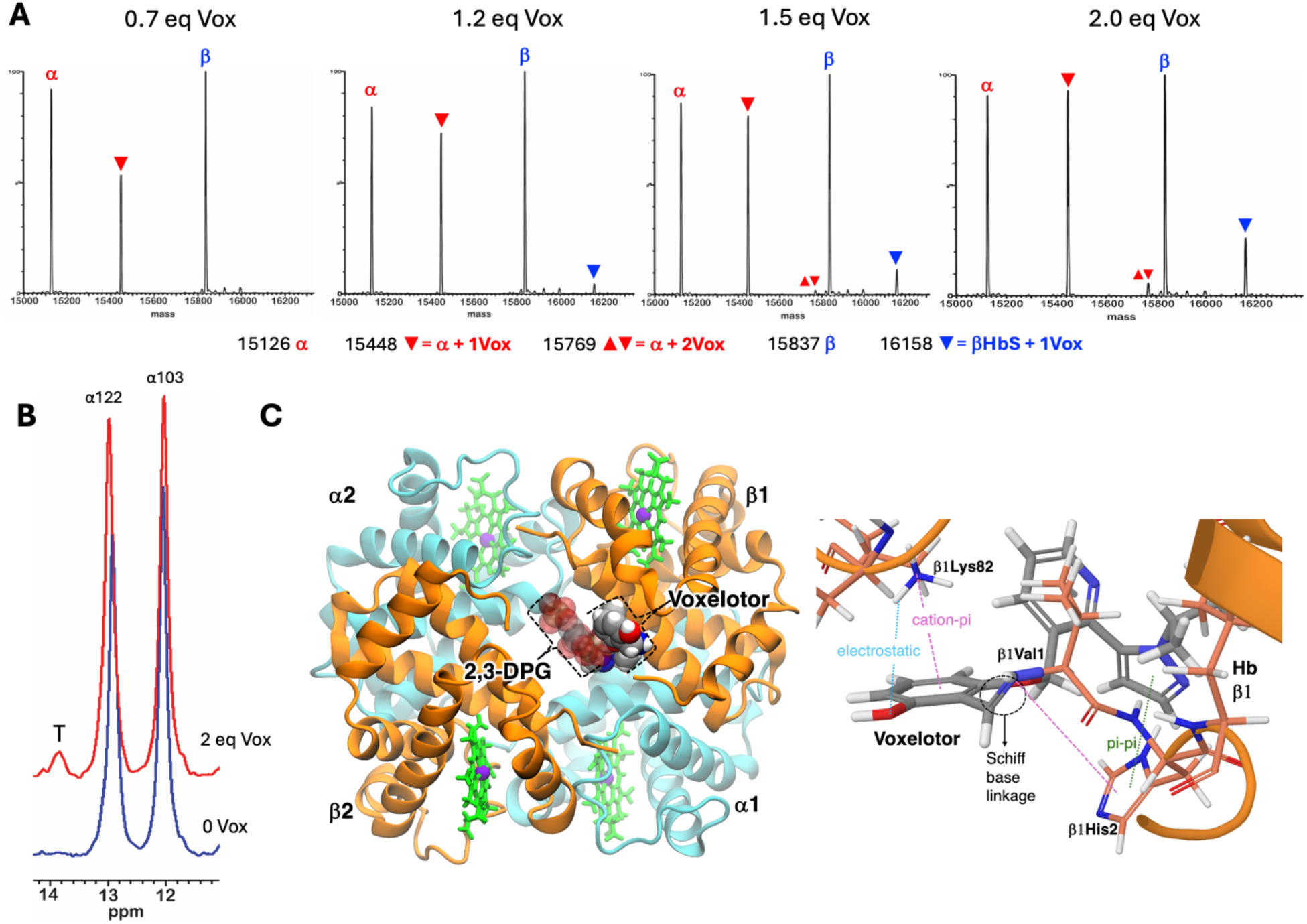
**A)** Lysate from a blood sample containing homozygous sickle cell hemoglobin (HbSS) mixed with increasing equivalents of Voxelotor (Vox). LC/MS was performed after trapping the Schiff base with 100 mM sodium borohydride (NaBH_4_); equivalents of Voxelotor added are shown above each spectrum compared to Hb concentration used (760 *μ*M). Comparison to previously reported binding studies under identical RBC lysis conditions^10^, shows that the stronger reductant used here allowed us to determine increased affinity and new secondary binding sites. **B)** 800MHz proton 1D spectra of 350 *μ*M HbSSCO lysate mixed with 700 *μ*M Voxelotor. The T-marker αTyr42 (H^η^) signal at 13.81 ppm is indicated by the letter T (see **Figure 2A**). αHis103 and αHis122 are N^ε2^H signals, two-proton intensity each (two α subunits per tetramer). **C)** Computational model of Voxelotor binding to T-state Hb at β1 N-terminus. (Right) The modeled Voxelotor (solid van der Waals surface) binding site overlaps with 2,3-DPG (transparent van der Waals surface) binding site. (Left) Voxelotor binding interactions modeled: (a) Schiff base linkage (1.3 Å) to Hb β1Val1; (b) cation-pi interaction (dashed magenta line) between the Voxelotor benzene ring and Hb β1Lys82 NH_3_^+^ group; and (c) pi-pi interaction between the pyrazole rings of Voxelotor and Hb β1His2; and (d) weak electrostatic interaction (dashed blue line) between the Voxelotor-OH and β1Lys82 NH_3_^+^ groups. Hb β1His2 also forms cation-pi interaction with Schiff base N^+^. The β N-terminus, His2, and Lys82 cations play a role in binding 2,3-DPG; our model shows that Voxelotor can interfere with the interactions of those groups involved in 2,3-DPG binding.

Voxelotor shows poor solubility in aqueous solutions, forming large, aggregated particles (**Figure S2**). Addition of drug to Hb results in turbid solutions. This raised concerns that the concentration of drug added was an unreliable metric of drug bound to Hb. Consequently, we determined the stoichiometry of binding for each of the NMR samples described below using LC/MS (**Figures 2C, 3B, S3, and S4**).

**Figure 2.**
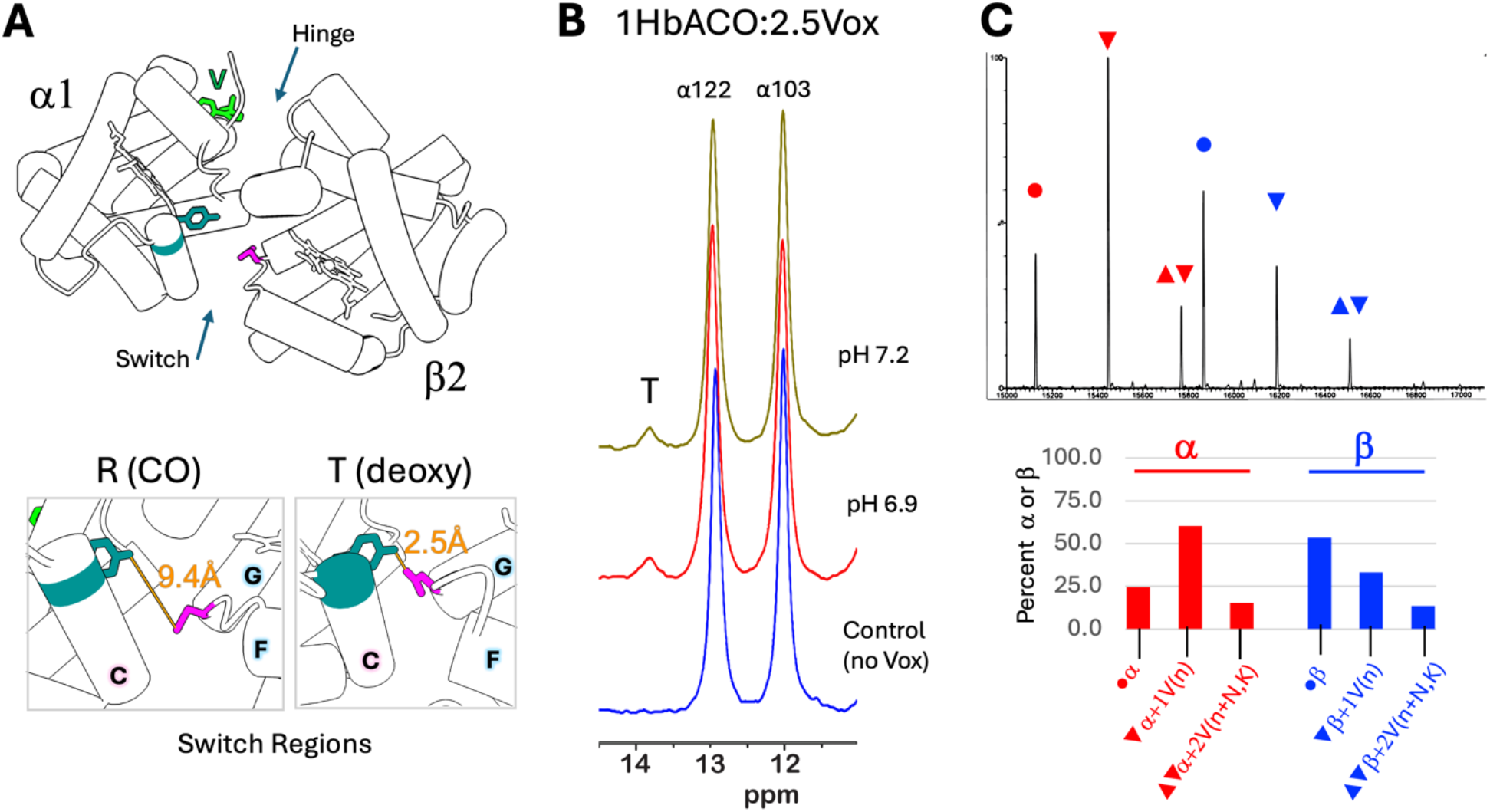
NMR 1D spectra and MS analysis of HbACO with Voxelotor. **A)** Crystallographic structure of the HbSSCO α1/β2 interface with Voxelotor bound; αTyr42 is teal; βAsp99 is magenta; Voxelotor is green (5e83.pdb). Boxes show the switch regions in the structures of HbSSCO (5e83.pdb) and deoxy HbA (1b86.pdb). Distances between H-bonding groups are shown. **B)** 800MHz proton 1D spectra of 200 *μ*M HbACO mixed with 500 *μ*M Voxelotor, pH 6.9 and 7.2 (almost identical to the concentrations used to prepare crystals: 186 *μ*M HbACO, 465 *μ*M Voxelotor, pH 7.4^1^). The T-marker αTyr42 (H^η^) signal at 13.81 ppm is indicated by the letter T; this signal is only present when there is a hydrogen bond between αTyr42 and αAsp99 in the switch region. αHis103 and αHis122 are N^ε2^H signals, two-proton intensity each (two α subunits per tetramer). **C)** NMR samples were assessed by LC/MS; the molar ratio of HbCO:Voxelotor (Vox) in panel **B** was estimated from MS intensities at mass: 15126 (α), 15448 (α+1Vox), 15769 (α+2Vox), 15867 (β), 16188 (β+1Vox), 16510 (β+2Vox). Percent occupancy is calculated from the intensity of each MS peak divided by the total intensities for that subunit (α or β). Since there are two α and β subunits, 50% would indicate one α or β subunit is fully occupied by one drug. Subunit +1V(n) correlates with binding predominantly to the N-terminus of that subunit; +2V(n+K) includes additional binding to side chains (Asn and Lys).

### NMR Shows Evidence of the T-Conformation Upon Drug Binding

We used NMR proton 1D and 2D NOESY spectra of Hb to assess the influence of drug on the resolved NMR signals. NMR signals of carbon monoxide bound Hb (HbSSCO or HbACO) are visible in the downfield region (11-14 ppm) and are indicators of quaternary structure. These peaks arise from buried sidechain-sidechain hydrogen bonding across α*/*β dimer interfaces. In **Figures 1B and 2B** peaks labeled α103 and α122 are from interactions across the α1/β1 interface; these do not change when the conformation shifts from R (oxy) to T (deoxy)^14,15^. We find that peak positions and patterns of spatial interactions for the 103 and 122 signals and valines at helix position E11 (NOEs, **Figure S3**) are unaffected by Voxelotor binding to HbACO; thus, drug binding does not appear to influence these local α1/β1 regions.

The most significant structural change between R and T conformations occurs in the switch and hinge regions of the α1/β2 (or by symmetry, α2/β1) interface (**Figure 2A**). An NMR peak at 13.81 ppm has been identified as a T-marker^16^ since this signal appears in the deoxy T conformation upon hydrogen bond formation between αTyr42 and βAsp99 (**Figure 2A**, switch region). Unexpectedly, this T-marker signal appears upon addition of Voxelotor to HbSSCO (**Figure 1B**) and HbACO (**Figure 2B, 2C**) under conditions of saturating carbon monoxide ligand. Voxelotor NMR signals do not resonate this far downfield^17^. The appearance of the T-marker signal correlates with bound drug and is not influenced by pH (**Figure 2B**).

Voxelotor binding to Hb has been shown to increase oxygen affinity. We also studied the NMR spectra of HbAO_2_ alone and in the presence of increasing amounts of Voxelotor (**Figure 3A**). Here we find similar results compared to the HbCO samples: The T-marker signal at 13.81 ppm appears upon addition of drug and the signal intensity of increases with drug concentration without effecting E11 signals that are sensitive to oxygen binding. The T-marker signal is apparent at low drug concentrations, where we observed only primary (α) site binding by LC/MS (**Figure 3B**). Thus, the R/T hybrid conformation is a property of Voxelotor binding to the α N-terminus.

**Figure 3.**
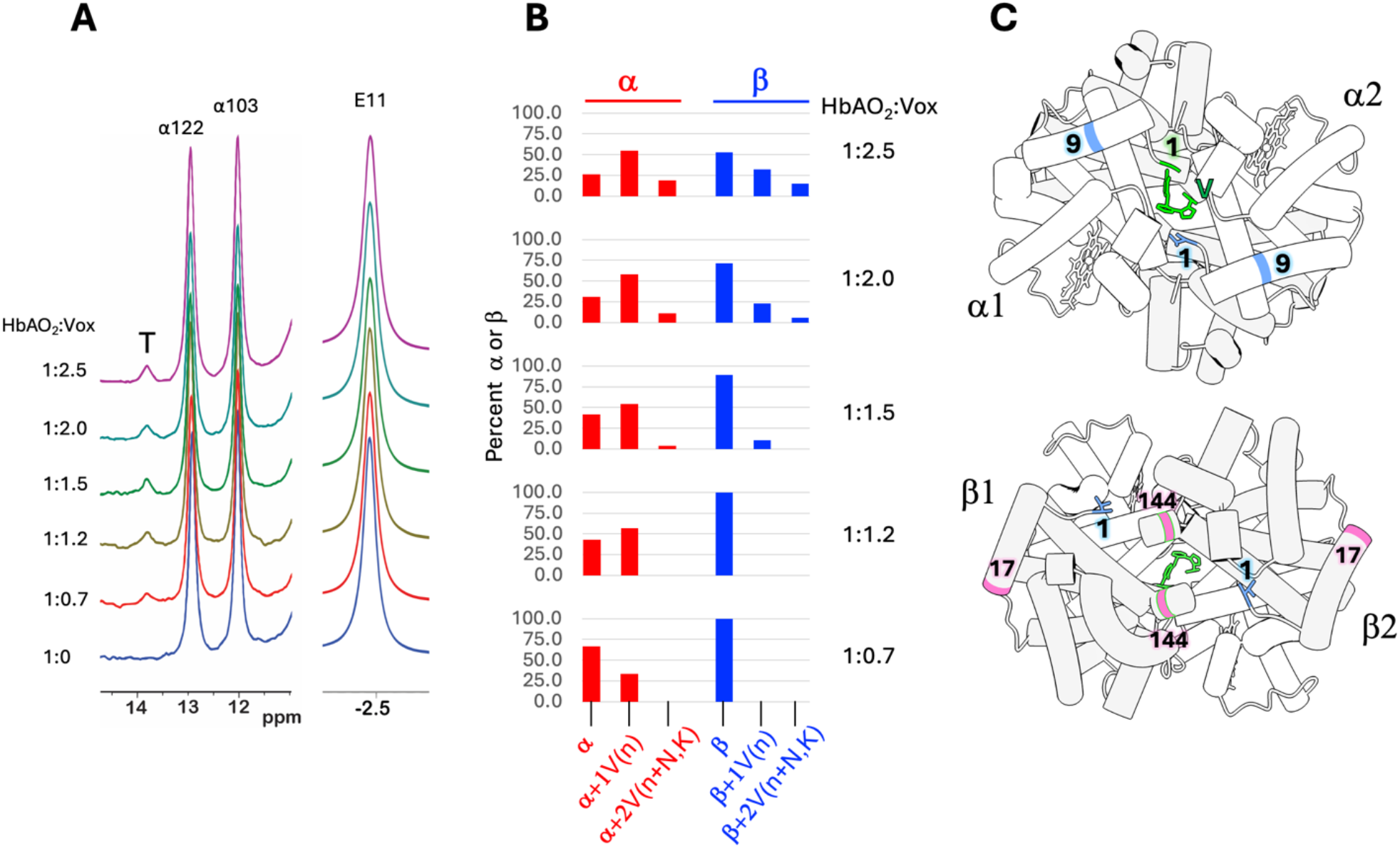
NMR 1D spectra and MS analysis of HbAO_2_ with Voxelotor. **A)** 800MHz proton 1D spectra of 200 *μ*M HbAO_2_ (pH 7.6, 5 mM sodium phosphate) with increasing Voxelotor concentration. The T-marker αTyr42 (H^η^) signal at 13.81 ppm is indicated by the letter T, positioned in the α1/β2 (or by symmetry, α2/β1) interface. αHis103 and αHis122 are N^ε2^H signals, two-proton intensity each (two α subunits per tetramer) at the α1/β1 (or α2/β2) interface. E11 marks αVal62 and βVal67 methyl signals sensitive to oxygen binding. **B)** NMR samples were assessed by LC/MS as in **Figure 2**. Ratios of tetramer Hb:Voxelotor were determined from MS intensities shown in **Figure S4**. Percent occupancy calculation and labeling are the same as in **Figure 2**. Each tetramer has two α and two β subunits; 50% binding indicates that one of the two subunits of that type is fully occupied. The N-terminus of one α subunit is the primary binding site. Once a single Voxelotor occupies the primary site, the second α N-terminus site is occluded and does not react with drug, but other secondary sites do become populated. The T-marker NMR signal is apparent at low drug concentrations where Voxelotor is bound at the primary site but not additional sites. Consequently, Voxelotor binding to the primary α N-terminus site is sufficient to drive a hybrid HbAO_2_ structure that includes a T-state feature. **C)** Binding sites mapped to the Hb-Voxelotor tetramer structure (5e83) based on tryptic digest significance (**Figure S5**). Top view, α subunits (white): α1Val1 strong (green); α2Val1, αAsn9 medium (blue). Bottom view, β subunits (grey): βVal1 medium (blue); βLys17, βLys144 medium-low (pink). Voxelotor bound to an α subunit is seen through the internal cavity the β subunit view.

To determine Voxelotor binding sites comprehensively, we performed tryptic digests on HbAO_2_ NMR samples alone and with two equivalents of drug bound per tetramer. nanoLC MS/MS detections suggested an ordered series of binding (starting with strongest) as 1) N-terminus of the α subunit (primary); 2) the N-terminus of the β subunit (**Figures 3C and S5**); 3) αAsn9, and weaker binding at 4) βLys17 and βLys113.

### Previous Studies Showed a Constrained Hb Structure and Underreported Drug Binding

NMR spectra are collected with the protein in solution, and since these spectra provide high-resolution information, they can reveal structural features not apparent when the protein is locked into a crystalline lattice. Here we see that Hb adopts a hybrid R/T structure when Voxelotor binds at the expected α site, even in the presence of a strong R ligand (CO).

Previous binding stoichiometry results (one drug per tetramer) were based entirely on interpretation of the crystal structure^1,17^. Published studies have been performed with solutions that contained substantial drug aggregation and the use of a weak reducing agent for detection have both contributed to an underreporting of bound drug^10^. However, both the α and β N-termini are known to be chemically reactive^18^. Our results show binding at the β N-terminus occurs near a 1:1 Hb(tetramer):Voxelotor stoichiometry. Progressive covalent modification of the β N-terminus (modeled in **Figure 1C**) could occur with the recommended dosage of 1.5 g/day over time and in individuals with high uptake. The consequences of binding to the β site have not yet been reported but might affect oxygen affinity and cooperativity^18^. It is likely that Voxelotor bound to the β N-terminus would interfere with 2,3-diphosphoglycerate (2,3-DPG) binding (modeled in **Figure 1C**) and could impact the delivery of oxygen to peripheral tissue. Depletion of this metabolite is known to have anti-sickling effects^19^. Inhibition of 2,3-DPG could create an illusion of successful treatment by increasing oxygen affinity and limiting sickling while causing adverse consequences. We hope that this new Voxelotor information will provide insight and inform future studies of similar compounds.

## Acknowledgements

We wish to thank the UCI Mass Spectrometry Facility, Felix Grun and Ben Katz for assistance with collection of protein mass spectrometry data and advice. Data were collected on a Waters Acquity UPLC Xevo G2-XS QTOF system (NIH supplemental funding support received by: J.S. Nowick (NIGMS GM097562), Vy Y. Duong (NIHGMS GM105938) and O. Cinquin (NIGMS GM102635).

We are grateful to Dmitry Fishman and Evan Garcia for advice and training on the Malvern DLS instrument.

This work was supported by National Institutes of Health instrumentation grant S10 OD016328 (P.D.G.).

## Authorship Contributions

MdcT wrote manuscript; performed research; collected data; analyzed data

EFC performed research; collected data; analyzed data

JS wrote manuscript; performed research; collected data

PG designed research; wrote manuscript; analyzed data

PTR designed research; wrote manuscript; performed research; collected data; analyzed data

MJC designed research; wrote manuscript; performed research; collected data; analyzed data

