## Supplemental Information for "Voxelotor (Oxbryta) Binds Multiple Hemoglobin Sites and Influences Protein Structure"

**Supplemental Methods**

### Protein and Drug

Voxelotor (aka Oxbryta or GBT440) was purchased from MedKoo (#329516) or MedChem Express (#HY-18681). Stock solutions of 100 mM drug in DMSO (SigmaAldrich) or d-DMSO (Cambridge Isotopes) were prepared (67.4 mg Voxelotor in 2 ml DMSO) and used fresh or aliquoted and stored at -80 °C.

Hemoglobin protein was purified from human blood donated for research and provided by the UCI Experimental Tissue Resource (samples deidentified except for sickle cell status, IRB HS #2012-8716 UCI-12-11). Erythrocytes were washed with phosphate-buffer saline (PBS, pH 7.2); sedimented cells were lysed with 3 volumes of water, rotating for 20 minutes at room temperature.

HbSS lysate was clarified with 10X PBS rotating for 10 min, then centrifugation at 14,000 g for 20 minutes, followed by filtration using a 0.22 μm filter. The concentration of HbSS in lysate (760 μM) was determined from the absorbances at 414, 541, and 575 nm[1]. HbSS lysate was tested for Voxelotor binding using LC/MS as described[2].

HbA lysates were stabilized in the CO-bound state by purging lysate with CO gas. Lysate was clarified by the addition of 10X PBS, rotating for 10 min, then centrifugation at 14,000 g for 20 minutes, followed by filtration using a glass prefilter (Whatman, Grade D). Removal of metHb was confirmed using visible spectroscopy[1] and NMR spectra measured with a large spectral width (120 ppm) to detect paramagnetic metHb signals.

The purification method for HbA was adapted from Sun and Palmer[3] with the following changes: The ion exchange column used was a Cytiva HiPrep Q-FF. Lysate was dialyzed against 20 mM Tris-HCl, pH 8.3 and eluted with a linear gradient to Tris-HCl, pH 8.3, 0.2 M NaCl. Protein purity was confirmed using SDS PAGE and LC/MS. Protein was dialyzed against 5 mM sodium phosphate, pH 7.6 (storage buffer), bubbled with CO gas and concentrated to 1 mM using a 10K MWCO centrifugation filter. Aliquots were frozen and stored at -80 °C.

Conversion of HbACO to $HbAO_2$ was accomplished by exposing the solution to strong visible light[4] (ARRI 650W Tungsten Spotlight) while purging the airspace with medical grade oxygen. The sample was rotated in contact with ice opposite the light source since the lamp emitted heat. Conversion to $HbAO_2$ was confirmed by quantitative deoxygenation using ascorbate (measuring the absorbance at 575 nm before and after addition of ascorbate) and NMR spectroscopy (the E11 signal is detected at -1.84 ppm for HbACO and at -2.45 for $HbAO_2$). Protein concentrations were determined using extinction coefficients of Meng and Alayash[1].

### NMR Spectroscopy

Voxelotor (diameter ~ 0.7 nm) is not soluble in aqueous solutions and can form large aggregates (diameter 0.4 to > 10 μm, **Figure S2**). We found that Hb samples became turbid upon addition of drug. Aggregation of drug at high concentrations appears to be a kinetic trap that inhibits binding. Our strategy to optimize binding was to use a low ionic strength buffer and, since some drug remained insoluble, we used LC/MS to quantitate the amount of drug bound to each Hb NMR sample immediately following NMR experiments. The ratios of protein to drug bound shown in **Figures 2 and 3** were determined from LC/MS (see below). The concentration of Voxelotor added to $HbAO_2$ compared to the quantity of Voxelotor bound determined by LC/MS is shown in **Figure S2**.

Voxelotor was mixed with protein and incubated overnight at 4 °C in the dark for each NMR sample. We tested the effect of ionic strength by adding sodium phosphate after drug binding. No change in the 1D NMR spectrum of HbACO-Voxelotor was seen when comparing spectra of 5 mM to 100 mM sodium phosphate solutions.

An NMR sample was prepared by diluting HbSS lysate 1:1 in water saturated with CO gas and $D_2O$ to make 350 μM HbSSCO. The space above the sample was flushed with CO gas for an additional 15 minutes prior to loading into an NMR tube and a 1D reference NMR spectrum was measured. Voxelotor was then added to a final concentration of 700 μM and another 1D NMR spectrum collected.

All HbA NMR samples were prepared from purified protein. HbACO samples for 1D measurements were prepared by mixing 1 mM HbACO, pH 7.2 in 5 mM sodium phosphate with a stock solution of 100 mM Voxelotor in d-DMSO to produce a final Voxelotor concentration of 5 mM drug. The sample was allowed to incubate overnight, split into two portions and then diluted to 200 μM HbACO with 100 mM sodium phosphate at pH 7.2 or 6.9. HbACO samples (300 μM) for 2D NOESY measurements were prepared in 5 mM potassium phosphate, pH 7.2 and incubated with 1500 μM Voxelotor overnight.

$HbAO_2$ will autoxidize to metHb near physiological pH and is usually studied at pH 8[5]. Using visible spectroscopy to detect conversion to metHb, we found that the oxygenated protein was stable at pH 7.6 for the drug incubation time (overnight) and NMR measurements. $HbAO_2$ (200 μM) samples were prepared in 5 mM potassium phosphate buffer, pH 7.6. 10% $D_2O$ was used as a lock frequency reference. Separate samples were prepared for each drug concentration from the same stock $HbAO_2$ solution. Final drug concentrations for each sample were 140, 240, 480, 800, 1600 μM. Additional d-DMSO was added to maintain a final d-DMSO concentration of 1.2% for all samples.

All NMR data were collected at 25 °C on a Bruker Neo 800 MHz NMR spectrometer equipped with an HCN probe. One-dimensional proton NMR spectra were collected with water suppression by gradient tailored echo (WATERGATE)[6] and using a spectral width of 25 ppm and 4K complex points. Two-dimensional NOESY experiments (HbACO) employed a mixing time of 100 ms and WATERGATE. A reference NOESY spectrum was collected on a 300 μM HbACO sample, then Voxelotor in d-DMSO was added to that same sample to a final concentration of 1800 μM drug. Another NOESY experiment was performed with identical acquisition parameters to the reference spectra: 25 ppm spectral width, 2048 x 128 complex points, relaxation delay = 1.5 sec. Spectra were processed using Bruker Topspin (4.3) and NMRfx Analyst (11.4)[7] software programs.

Processing parameters were identical for each sample: 10-Hz line broadening was applied to 1D spectra; 2D spectra were processed with a squared sine bell shifted 0.3 window and zero-filled to double the size in the indirect dimension. A low-frequency solvent filter was applied, and all spectra were baseline corrected using polynomial functions.

#### Dynamic Light Scattering

Since the human eye cannot see particles less than 10 μm in size, we used Dynamic Light Scattering (DLS) to determine whether aqueous solutions were monodispersed or contained drug aggregates (polydisperse). Voxelotor solutions were prepared from 10 mM drug in DMSO diluted into PBS, pH 7.2 to final concentrations of 100 μM and 300 μM. Samples were prepared and vortexed immediately prior to measurements. DLS experiments were performed on a Malvern Zetasizer Nano ZS using the following standard Malvern parameters: Dispersant was set to PBS; measurement was set to 173 backscatter, Automatic Measurement Duration was selected with 3 records taken, each an average of 18-22 measurements per record. Selected cell was set to Quartz Cuvette, Zen2112. The temperature was set to 25 °C, with a 5-minute equilibration time. Polydispersity was observed at concentrations above 300 μM and correlated with a decrease in bound drug detected (**Figure S2**).

#### LC-MS method for analysis of protein conjugation

Since α and β subunits are detected with different efficiency by MS, subunits were assessed individually by adding the total intensities of drug bound peaks for each subunit and then dividing by the total intensity for that subunit. The final ratio was the sum of the percentage bound values for each subunit type (α or β).

*Trapping the Schiff base:* A fresh solution of 1 M sodium borohydride (#S0480, Tokyo Chem Ind) was prepared in ice-cold $H_2O$ immediately prior to use. 10 μl of 1M sodium borohydride was added to 100 μl of 100 μM Hb-Voxelotor to a final concentration of 100 mM sodium borohydride. This was allowed to react on ice for 30 minutes.

*Comparison of Reducing Agents:* Human metHb (SigmaAldrich, #H7379) was used to test the efficiency of Schiff base reduction using sodium borohydride compared to sodium cyanoborohydride (Oakwood Chemical, #044871). 100 μM metHb was prepared in 100 mM sodium phosphate, pH 7.1. Voxelotor was added to a final concentration of 100 μM. The mixture was allowed to incubate for one hour at room temperature and split to three equal portions. 1 M sodium borohydride and 1 M sodium cyanoborohydride were prepared fresh with cold $H_2O$. 1 M reductant solutions were diluted into metHb samples to yield final reductant concentrations of 50 mM (both cyanoborohydride and borohydride) and 100 mM borohydride. Samples were incubated on ice for 30 minutes.

All samples were diluted to 2-10 μM Hb concentration for LC/MS measurements. Protein conjugation to Voxelotor was measured by LC/MS analysis of the intact protein sample (ACQUITY UPLC H-class system, Xevo G2-XS QTOF, Waters). Proteins were separated from reaction buffer salts using a phenyl guard column at 45 °C (ACQUITY UPLC BEH Phenyl VanGuard Pre-column, 130Å, 1.7 µm, 2.1 mm X 5 mm, Waters). A 5-minute method (**Table S.1**) used a 0.2 mL/min flow rate of a gradient of Buffer A consisting of 0.1% Formic Acid in water (Water LC-MS #9831-02, J.T. Baker; Formic Acid LC-MS #85178, Thermo Scientific) and Buffer B, Acetonitrile (Acetonitrile UHPLC/MS #A956, Thermo Scientific).

**Table S1.** Protein Method Gradient

| Time (mins) | Flow (mL/min) | %A | %B | Curve |
|---|---|---|---|---|
| Initial | 0.2 | 100 | 0 | 6 |
| 0.5 | 0.2 | 100 | 0 | 6 |
| 2 | 0.2 | 10 | 90 | 6 |
| 2.5 | 0.2 | 10 | 90 | 6 |
| 4 | 0.2 | 100 | 0 | 6 |
| 5 | 0.2 | 100 | 0 | 6 |

The Xevo Z-spray source was operated in positive ion MS resolution mode, with a capillary voltage of 3000 V and a cone voltage of 40 V (NaCsI calibration, Leu-enkephalin lock-mass). Nitrogen was used as the desolvation gas at 350 °C and a total flow of 800 L/hr. Spectra were acquired in the 400 - 4000 Da range. Total average mass spectra were reconstructed from the charge state ion series using the MaxEnt1 algorithm from Waters MassLynx software V4.1 SCN949 according to the manufacturer's instructions. To obtain the ion series described, the major peak of the chromatogram was selected for integration before further analysis. Spectra were processed with identical parameters (chromatogram peak selection from 2.5 – 3.5 minutes, and maximum entropy).

#### nanoLC-MS/MS

25 µg of human $HbO_2$:Vox 1:2 solution (NMR sample, pH 7.6, reduced with 100 mM sodium borohydride) was rendered in 8 M in urea, 10 mM in tris(2-carboxyethyl)phosphine (TCEP) and 0.1 M in triethylammonium bicarbonate (TEAB), pH 8.0 then incubated for 30 minutes at 37 °C. The resulting samples were diluted with 0.1 M TEAB (pH 8.0) to 6 M urea then supplemented with 0.25 µg of recombinant LysC protease. After overnight (~20 h) incubation at 37 °C, samples were further diluted to 1 M urea with 0.1 M TEAB (pH 8.0) then supplemented with 0.25 µg trypsin followed by overnight (~20 h) incubation at 37 °C. Samples were then acidified with formic acid to 3% (v/v) final concentration and pH < 3.0 followed by peptide desalting using C18–SCX stacked StageTips as described[8]. After drying the eluate under vacuum, desalted peptides were redissolved in 0.1% formic acid in water.

75 micron inside diameter x 25 cm nanocapillary columns were packed in-house with ReproSil-Pur C18-AQ beads (1.9 µm diameter; Dr. Maisch GmbH). Using an EASY-nLC 1200, columns were equilibrated with 0.1% formic acid (solvent A) in water followed by

sample injection and a discontinuous gradient from 0 to 5% solvent B (93% CH3CN in 0.1% formic acid/water) over 5 min then to 23% B over 60 min then to 35% B over 15 min, at a flow rate of 250 nL/min. Column eluate was electrosprayed from the column end into an LTQ Orbitrap Velos Pro mass spectrometer, acquiring FTMS precursor spectra at 60,000 resolution followed by MS2 spectra (FTMS, 7500 resolution) of the 15 most intense >+1 charged precursor ions with intensity > 2000, fragmenting by Higher-Energy Collisional Dissociation (30% Normalized Collisional Energy). After two fragmentations within 30 sec ions were dynamically excluded for 40 sec via a 500-entry list with early expiration from the list after a detection within the exclusion period that fell below S/N = 2.0.

In Mascot Server 2.8, after configuring custom drug modifications for adductation of lysine sidechain and protein N-terminus, raw data were searched (MS/MS ions; target/decoy) against the human proteome (UniProt) plus a custom database of common contaminants, with trypsin specificity and variable modifications including Oxidation (M), Deamidated (NQ) and custom modifications. Searches specified precursor and product mass tolerances of 20 ppm, allowing +2, +3, +4 charge states and one missed cleavage per peptide. Output data were thresholded at 2% peptide-level false discovery rate (FDR).

**Molecular Docking and Energy Minimization**

Computational modeling of Voxelotor-bound Hb structure was accomplished using Schrödinger software suite Release 2025-2[9]. Atomic coordinates of the Hb T-state were obtained from PDB ID 1B86 and isolated from the bound 2,3-DPG metabolite. Atomic coordinates of Voxelotor were isolated from PDB ID 5E83. Both coordinates were then structurally optimized for atomistic modeling in terms of adding missing atoms (such as all H atoms), bonds, and assigning accurate charges through the Schrödinger software suite's Maestro platform. Glide docking software[10] was then used to identify potential binding site of Voxelotor around the 2,3-DPG binding site. Voxelotor was successfully docked at the N-terminus of Hb $\beta$1 subunit. The docked ligand was next attached to b1Val1-NH2 terminus. The docked ligand was next attached to b1Val1-NH2 terminus. The drug and Hb residues at and near the drug binding site region underwent a local structural minimization with the OPLS_2005 force field[11]. An optimized Schiff base linkage of 1.3 Å was obtained.

Finally, restrained minimization of the Voxelotor-bound Hb structure was conducted using the OPLS_2005 force field[11]. VMD software[12] was then used to structurally align this optimized Voxelotor-bound Hb structure to the 2,3-DPG bound Hb (PDB ID:1B86) at the protein backbone of the 2,3-DPG binding site. 2,3-DPG binding site residues were assigned as those within 5 Å of the 2,3-DPG binding site.

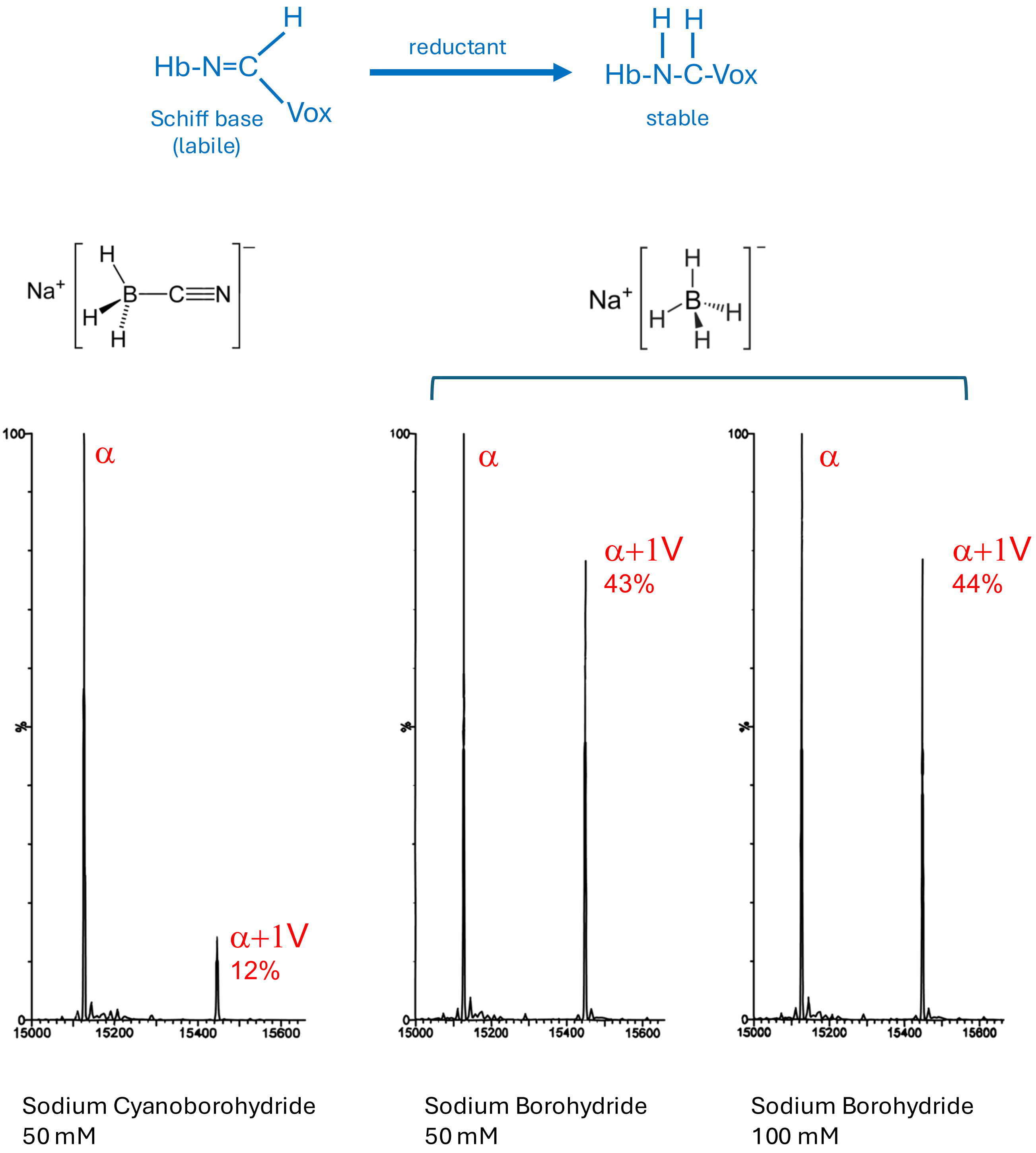


**Figure S1.** Reductant efficiency. One equivalent of Voxelotor were added to a 100 μM solution of metHb (Sigma). The sample was split and either sodium cyanoborohydride (50 mM) or sodium borohydride (50 mm or 100 mM) were added. LC/MS show that 3.6 times more bound Voxelotor is detected when the stronger reductant, sodium borohydride is used.

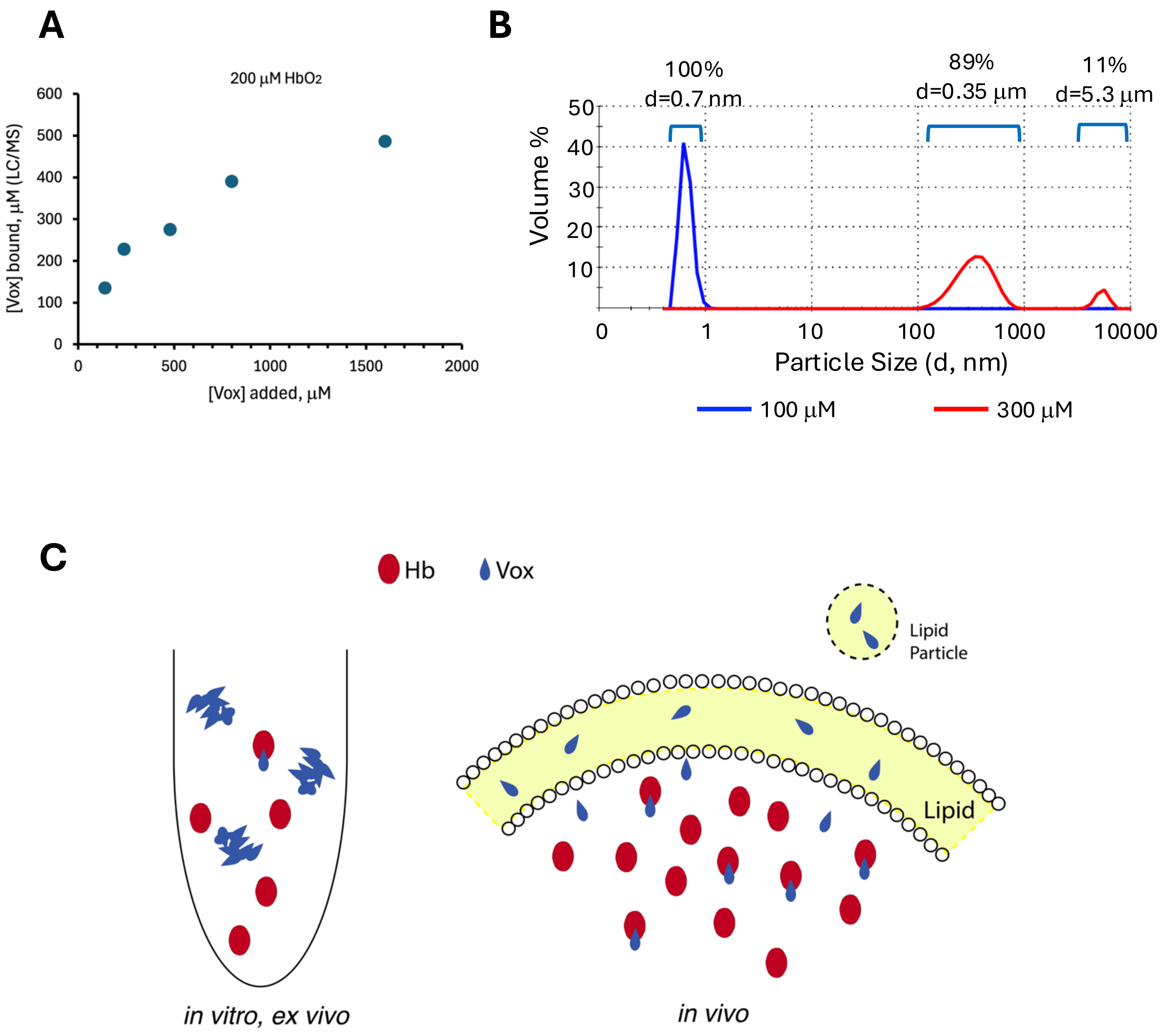


**Figure S2.** Voxelotor reactivity and aggregation in aqueous solutions measured by Dynamic Light Scattering (DLS). **A)** Comparison of the concentration of Voxelotor added vs the amount calculated to be bound to 200 μM $HbO_2$ based on LC/MS. **B)** DLS of Voxelotor in PBS: 100 μM (blue) shows the drug is monodisperse at this concentration; 300 μM (red) reveals large particle formation invisible to the eye. **C)** Voxelotor aggregation *in vitro/ex vivo* complicates interpretation of binding data since drug is less available and can lead to an underestimate of drug affinity. In the body, drug can be delivered in fat particles, then partition into the cell membrane which is a highly hydrophobic liquid phase. Slow transfer of drug into the cell inhibits drug aggregation, especially since the concentration of the binding target, Hb, is very high (~ 5mM). Once Voxelotor attaches to Hb in the cell, it cannot aggregate. Consequently, the effective concentration of Voxelotor is higher in the cell compared to *in vitro/ex vivo* measurements where it is kinetically trapped in insoluble particles.

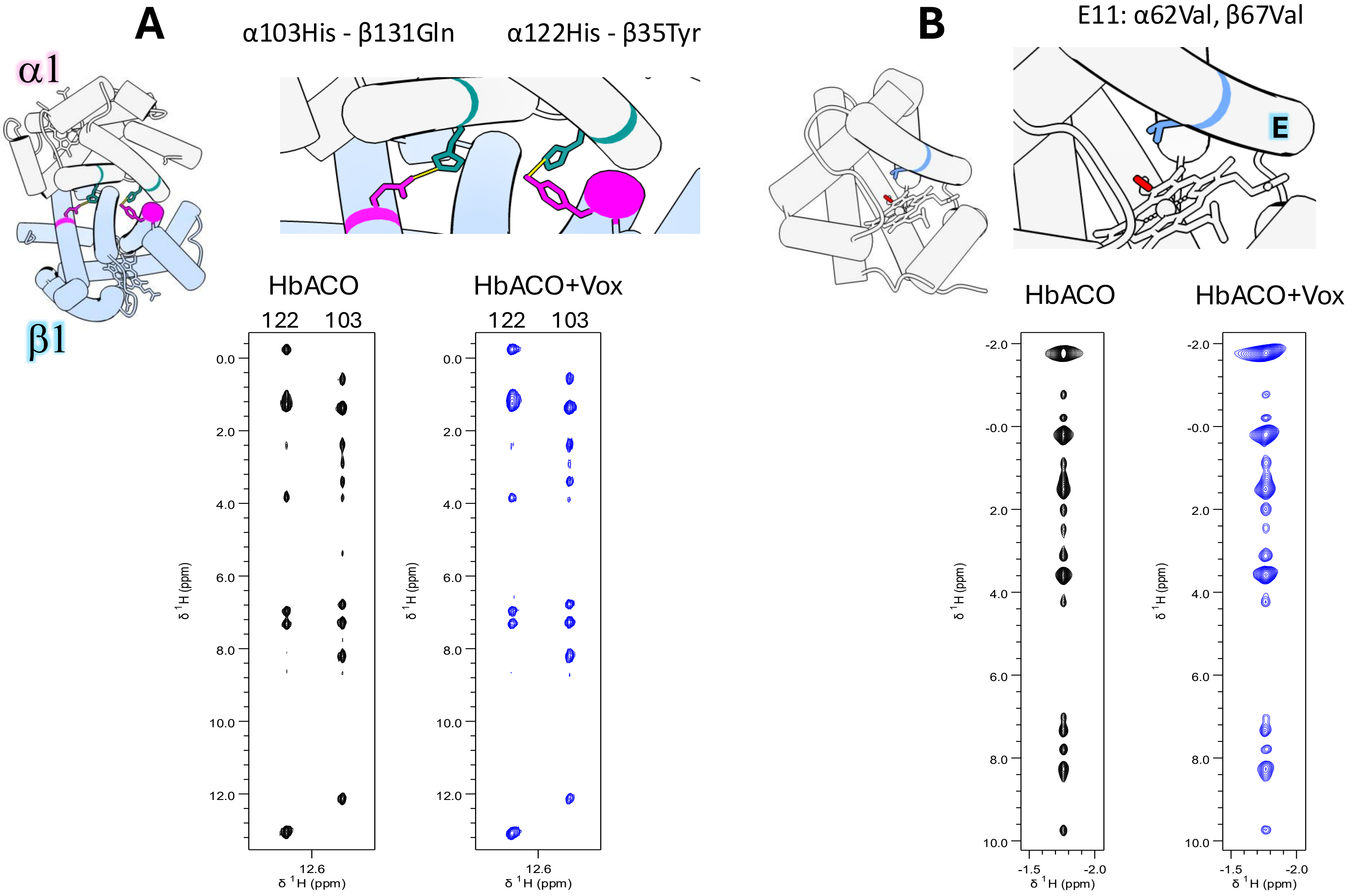


**Figure S3**. 2D NMR NOESY ($\tau_{mix}$ = 100 ms) sections of HbACO alone and with Voxelotor. **A)** Histidines 103 and 122 form key hydrogen bonds across the α1/β1 interface. **B)** Valines that occupy position 11 in the E helix have contact with both the heme and ligand. Chemical shifts and NOE patterns in both **A)** and **B)** are very similar indicating Voxelotor has minimal structural effect in these regions. Binding of Voxelotor does increase linewidths, consistent with an increase in protein dynamics.

**Figure S4.** $HbAO_2$ with increasing Voxelotor concentration. Aliquots of NMR samples (**Figure 3**) were diluted into 100 mM sodium borohydride to trap the Schiff base.

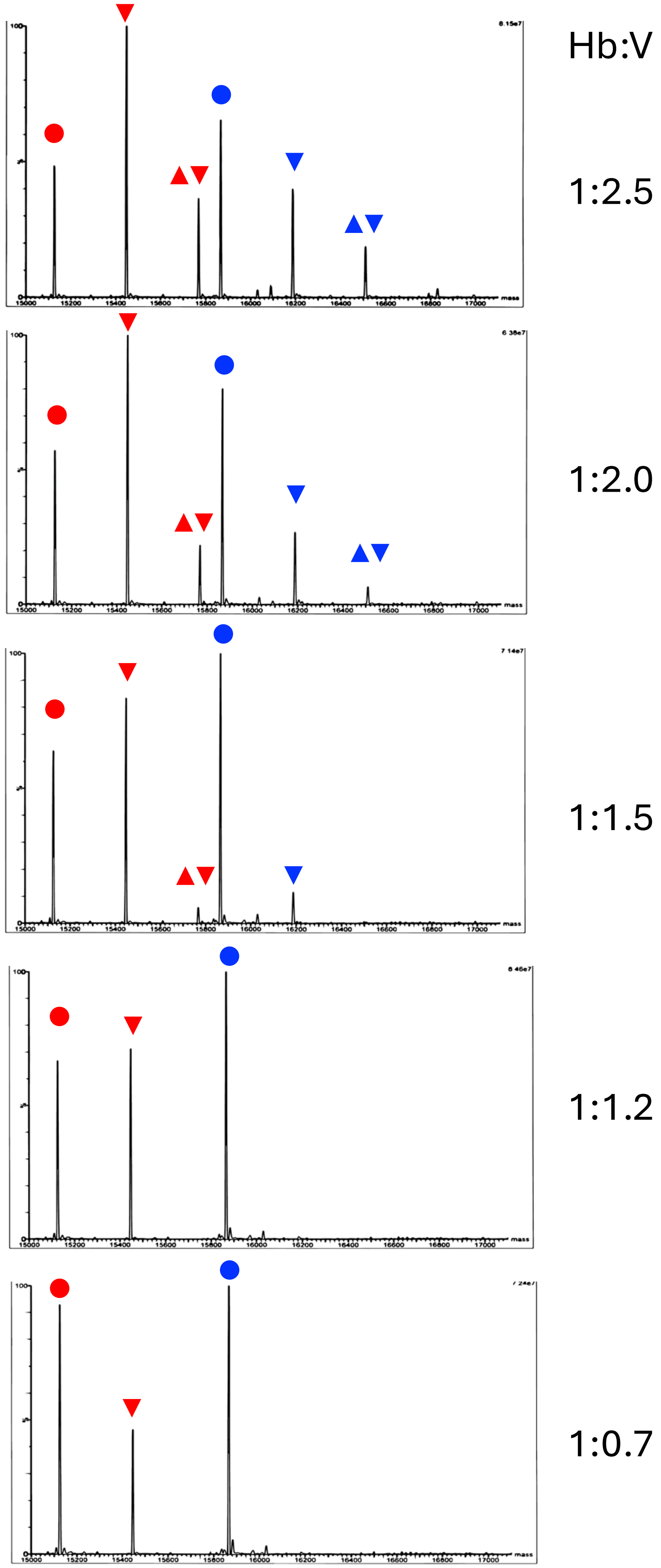


| mass | | |
|---|---|---|
| 15126 | ● | α |
| 15488 | ▼ | α+1v |
| 15769 | ▲▼ | α+2v |
| 15867 | ● | β |
| 16118 | ▼ | β+1v |
| 16510 | ▲▼ | β+2v |

**(a)**

MVLSPADKTNVKAAWGKVGAHAGEYGAEALERMFLSFPTTKTYFPHFDLSHGSAQVKGHGKKVADALTNAVAHVDDMPNALSALSDLHAHKLRVDPVNFKLLSHCLLVTLAAHLPAEFTPAVHASLDKFLASVSTVLTSKYR

| | | |
|---|---|---|
| **n**-VLSPADK | 2 | 0.0006 |
| **n**-VLSPADKTNVK | 20 | 2.3E-07 |
| **n**-VLSPADKT**N**VK | 5 | 6E-06 |
| AAWG**K**VGAHAGEYGAEALER | 2 | 0.0014 |

**(b)**

MVHLTPEEKSAVTALWGKVNVDEVGGEALGRLLVVYPWTQRFFESFGDLSTPDAVMGNPKVKAHGKKVLGAFSDGLAHLDNLKGTFATLSELHCDKLHVDPENFRLLGNVLVCVLAHHFGKEFTPPVQAAYQKVVAGVANALAHKYH

| | | |
|---|---|---|
| **n**-MVHLTPEEK | 1 | 0.0034 |
| **n**-VHLTPEEK | 7 | 0.00093 |
| **n**-VHLTPEEKSAVTALWGK | 5 | 1.3E-05 |
| VHLTPEE**K**SAVTALWGK | 1 | 0.00035 |
| SAVTALWG**K**VNVDEVGGEALGR | 1 | 1.1E-05 |
| VVAGVANALAH**K** | 1 | 1.1E-05 |

**Fig. S5**. Voxelotor-labeled peptides detected in (**a**) Hb $\alpha$ and (**b**) Hb $\beta$, in tryptic digests of $HbAO_2$:Voxelotor = 1:2 molecules bound (**Figure 3).** The pH of this NMR sample was 7.6 to stabilize the oxygenated state. Lysine side chains (pKa ~ 10.5 if exposed) might be slightly more reactive at this pH compared to the range of physiological pH values; however, during prolonged exposure at 37 °C these positions in Hb might react with Voxelotor. Beneath protein sequences are shown the tryptics with labeled position bold red (“n-“ denotes protein N-terminus either demethioninylated or Met-containing). Cyan: Deamidated Asn.

Numerical columns: (1) number of detections, (2) best Mascot expectation value (approximate probability of misidentification). Values considered highly significant are highlighted yellow.
